# Unprecedented urban freshwater microviral survey highlights shift from sewage to river water communities

**DOI:** 10.64898/2026.11.03.763685

**Authors:** A Langlais, C Dumont, J Mougin, S Rechdaoui-Guérin, C Roose-Amsaleg, F Enault, A Quaiser

## Abstract

Microviruses are abundant yet still poorly resolved components of virosphere, particularly across urban freshwater continua. Here, we investigated complete microvirus genomes recovered from sewage water (SW), treated wastewater (TW), and river water (RW) along an urban SW–TW–RW continuum, providing an extensive genomic dataset of 18,449 complete genomes grouped into 825 genera. This catalog enabled a high-resolution assessment of microvirus diversity, taxonomy, and ecological distribution across anthropized aquatic compartments. All currently described microvirus orders were detected, with *Gokushovirales* dominating the dataset and *Betagokushoviridae* emerging as the most prevalent family in TW and RW samples. In contrast, *Alpavirales* were enriched in SW but strongly depleted after treatment, consistent with their association with gut-related bacterial hosts. Alpha- and beta-diversity analyses showed that water type was the main driver of microvirus community structure, whereas seasonal effects were weak and not robust under the hydrological conditions of the sampling year. Wastewater treatment plants thus acted as strong biological filters, reshaping viral assemblages and selecting compartment-specific microvirus populations. Finally, the pronounced and reproducible shifts observed across the continuum suggest that selected microvirus groups may serve as promising bioindicators of anthropogenic influence in freshwater systems.

## 1. INTRODUCTION

Freshwater ecosystems provide essential ecological functions and ecosystem services, including nutrient cycling, organic matter decomposition, and water purification. However, these ecosystems are increasingly exposed to anthropogenic pressures, particularly in urbanized environments where human activities profoundly alter hydrological, physicochemical, and biological conditions (Carpenter *et al*., 2011). Urban rivers are subjected to multiple stressors, including nutrient enrichment, chemical contamination, stormwater runoff, habitat modification, and the discharge of domestic and industrial effluents. These cumulative pressures can disrupt ecosystem functioning and biodiversity, ultimately affecting water quality and ecological integrity. In recent decades, additional challenges such as climate change, population growth, and increasing urbanization have intensified these perturbations. Among the various sources of pollution, fecal contamination has emerged as a recurrent concern in urban waters, with potential consequences for both ecosystem and public health. Wastewater treatment plants (WWTPs) constitute a major interface between human activities and aquatic environments. Although designed to reduce pollutant loads before discharge, WWTP effluents remain important sources of nutrients, chemical contaminants, and microorganisms released into receiving waters (Ruprecht *et al*., 2021). Microbial communities are key components of both freshwater ecosystems and WWTPs, where they drive critical biogeochemical processes. Consequently, numerous studies have investigated the structure and dynamics of microbial assemblages in urban rivers and wastewater treatment systems (Price *et al*., 2018; Mansfeldt *et* al., 2020; Wang *et al*., 2022). Increasing evidence shows that WWTP discharges can cause local but measurable restructuring of riverine microbial communities, increasing the relative abundance of wastewater-associated and fecal indicator taxa, modifying alpha diversity and community composition near outfalls, and introducing microorganisms linked to nutrient cycling, sewage treatment processes, and potentially pathogenic lineages (Drury *et al*., 2013; Ruprecht *et al*., 2021). These effects may partially attenuate with distance downstream depending on dilution, hydrology, and recovery capacity of the receiving stream (Castelar *et al*., 2022).

The composition and dynamic of microbial communities are influenced by the presence of viruses (Suttle, 2007). Most of them are bacteriophages and consequently are involved in microbial communities’ control and biogeochemical cycles (Sime-Ngando, 2014; Elbehery and Deng, 2022). Understanding viral ecology, particularly viral diversity and dynamics is therefore essential to characterize ecosystem functioning at both local and global scales (Jacquet *et al*., 2010; Sommers *et al*., 2021). In anthropized aquatic environments, viral communities are shaped by the mixing of human-, animal-, and environment-associated microbiota, as well as by human-induced physicochemical pressures (Huang *et al*., 2024). Urban freshwater systems are therefore characterized by complex and diverse viral assemblages, including numerous bacteriophages originating from multiple ecological reservoirs (Chen *et al*., 2023; Pollegioni *et al*., 2022). Among anthropogenic inputs, wastewater represents a particularly important source of viruses, concentrating microbial and viral populations from human, animal, and industrial origins, including bacteriophages associated with the human gut microbiome (Ballesté *et al*., 2022). Previous viromics studies have shown that sewage viromes largely reflect human-associated viral communities, making them a concentrated signal of urban microbial activity (Adriaenssens *et al*., 2021; Niu *et al.,* 2026). In WWTPs, this signal is transformed, modifying the microbial and viral community, by exerting strong physicochemical and biological pressures (Guajardo-Leiva *et al*., 2025). High microbial densities within treatment systems provide favorable conditions for phage replication, while treatment steps can lead to the elimination, selection, or concentration of specific viral populations (Saguti *et al*., 2025). Consequently, treated water viromes can conserve a WWTP signature, which will be exported to the receiving environment (Niu *et al*., 2026). Rivers, as a receptor environment, are composed of diverse microbial and viral sources. Those sources can be environmental, as aquatic communities, soil communities from rainwater runoff or linked to the anthropized area, as urban or industrial effluent, such as WWTP outlet (Elbehery and Deng, 2022). Influence of the treated water on river viral communities was detected downstream of the discharge, where the community profile was partly shaped by hydraulic mixing with the effluent and gradually became less similar to it with distance (Niu *et al*., 2026).

Among these viral communities, microvirus form a group often described as ubiquitous, abundant or even dominant across diverse environments, including marine water, freshwater, soil and host-associated microbiomes (Krupovic and Forterre, 2011; Roux *et al.,* 2012; Quaiser *et al*., 2015; Kirchberger *et al*., 2022; Olo Ndela *et al*., 2023; Wu *et al*., 2024). Microviruses were recently reclassified into the *Microviricetes* class containing seven orders and 21 families (Black *et al*., 2026). These small phages, with a circular and single-stranded DNA genome of size ranging from 3 to 10 Kb, potentially infect many different bacteria and might have a broad impact on the microbial community. Interestingly, this viral group has been identified in WWTPs representing between 10 and 30 % of infectious somatic coliphages in raw and treated wastewater from four urban WWTPs (Bichet *et al*., 2024). In most studies, this viral group is treated as a whole, without distinguishing subgroups. Thus, their spatial dynamics at different taxonomic levels along the continuum from sewage to treated water and river ecosystems remain largely unexplored.

In this context, our study aims to fill several important knowledge gaps about microvirus community structure in highly anthropized environments. The Seine River represents a relevant model system to investigate these dynamics. As a major European river under strong anthropogenic pressure, particularly in the Paris area, it receives substantial urban and industrial inputs. While previous studies have characterized bacterial, archaeal, and eukaryotic communities in the Seine River (Bagagnan *et al*., 2024; Hervé *et al*., 2025), viral diversity remains underexplored and has primarily focused on pathogenic viruses detected by PCR approaches (Prevost *et al*., 2015; Wurtzer *et al*., 2020).

Here, we applied metagenomic sequencing approach to study microviral communities along the continuum from sewage water to treated water and river environments and focused particularly on the microviral diversity and dynamic. In addition, we explored potential host associations to better understand the ecological origins and distribution of these viruses across human-associated and environmental niches.

## 2. EXPERIMENTAL PROCEDURES

### 2.1. Sampling process

River water (RW) samples were collected from the Seine River at 3 locations: Choisy (48°45’59.0”N, 2°24’43.9”E), Ivry (48°48’52.7’’N, 2°24’37.0’’E) and Colombes (48°55’58.3’’N, 2°14’30.6’’E) and from the sewage water (SW) and treated water (TW) of 2 WWTP: Seine Valenton (SEV; 48°45’43.721’’N, 2°27’7.887’’E) and Seine Centre (SEC ; 48°55’55.443’’N, 2°14’45.385’’E) (Figure 1). Sampling sites are located in the Paris area, on a river section of 50 km between Choisy, upstream of Paris, and Colombes, downstream of Paris. Between those points, two tributaries flow into the Seine River: the Marne River and the Oise River. SEV and SEC are two urban WWTP, their catchment areas serve 3 million equivalent inhabitants (2M and 1M respectively). Sampling campaigns were carried out seasonally in March 2024, June 2024, September 2024 and January 2025.

**Figure 1.**
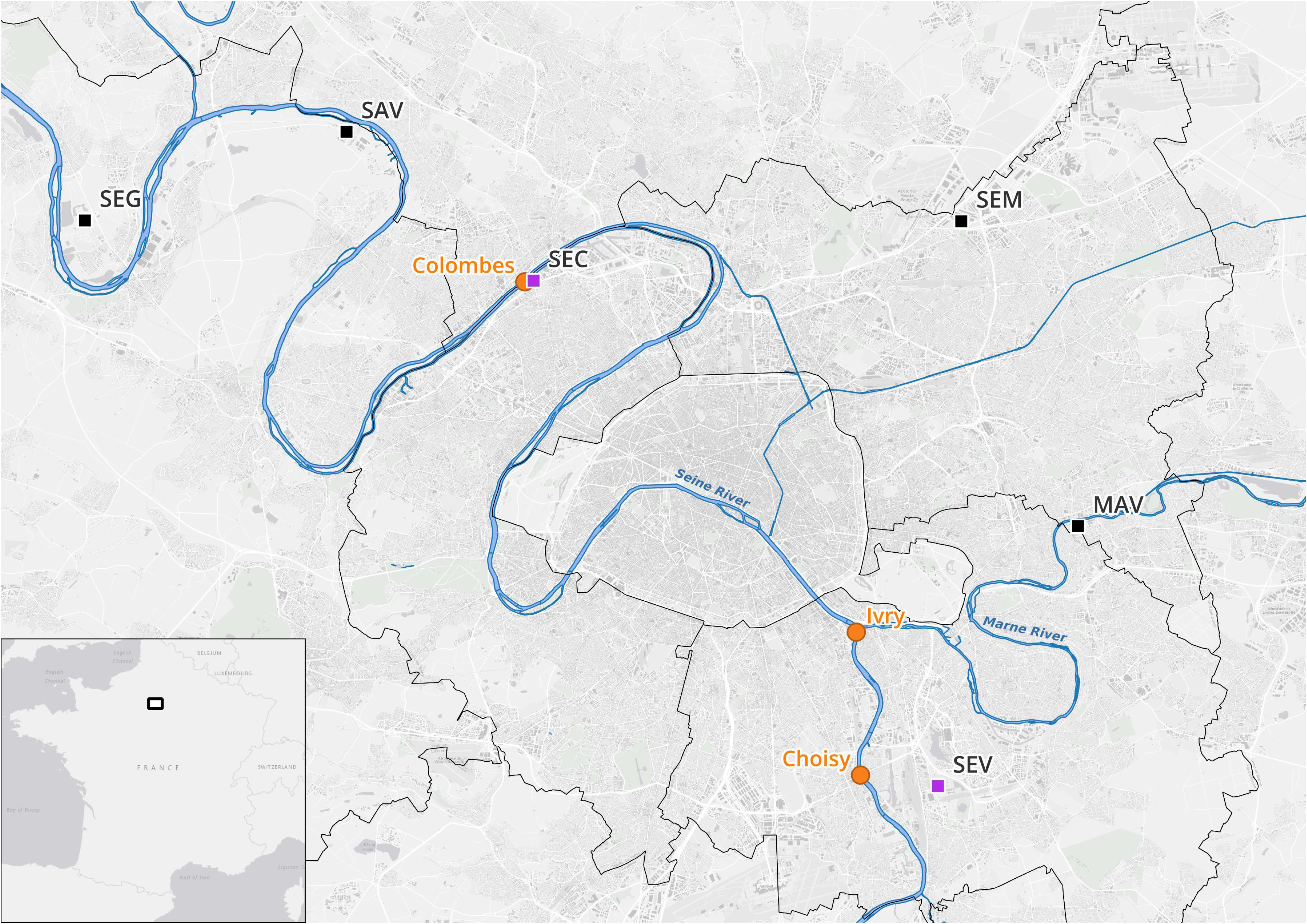
Location of sampling sites in the Paris region. Orange circles indicate the three river sampling points (Colombes, Ivry, Choisy). Purple squares represent the two sampled WWTP (SEC, SEV). Black squares correspond to other stations in the region discharging into the Seine or Marne rivers (SAV, SEG, SEM, MAV). The inset in the bottom left locates the study area (white rectangle) within France.

Water of the Seine River was sampled in triplicates by collecting 3 x 10 liters, with a buck, previously rinsed with Seine River water, near the water surface along the central axis of the river. Four liters of the SW from each WWTP were collected using auto-sampler devices, representing a 24-hour composite sample. We also sampled 15 L of the TW of both WWTP. All samples were stored at 4°C until processing the same day.

### 2.2. Sample preparation

Two different sample preparations were assessed. The SWs were centrifuged at 3500g for 15 minutes at 4°C to separate the free viruses from microorganisms and suspended matter. The supernatant was filtered through a 0.45µm membrane filter (Bottle-Top Vacuum Filtration Systems, PES, VWR). One-fifth of the pellet was conserved at -20°C for microbial metagenome study. The remaining pellet was resuspended in Potassium-Citrate buffer (pH7) (Williamson *et al*., 2003), incubated for 1h under vigorous movement at 20°C to detach viruses from the suspended matter. This suspension was centrifuged at 3500g for 15 min, and the supernatant was filtered through a 0.45µm membrane filter. The filtered suspensions from both were amended with 10% PEG8000 and 0.6% NaCl and incubated at 4°C during 7 days to precipitate viruses (Colombet *et al*., 2007).

Concerning the TW and the RW samples, a succession of filtrations was performed. Samples of 10 liters were filtered three times with increasingly finer filters: 1mm, 5µm and 0.45µm. Then a cross-flow filtration was performed on 30kDa filter columns (Rexeed 25A hemodialyzer, Hemotech) to concentrate viruses and to reduce the volume to less than 500mL. Viruses were precipitated with 10% PEG8000 and 0.6% NaCl and incubated during 7 days at 4°C.

After PEG precipitation, centrifugation at 8000g during 20 minutes was performed. Pellets containing viruses were resuspended with 1M KCl, the suspensions were concentrated with 100kDa ultrafiltration spin columns (Vivaspin 20, 100 kDa MWCO Polyethersulfone, Sartorius) to a final volume inferior to 6mL and stored at -20°C until nucleic acid extraction.

### 2.3. Nucleic acids extraction and sequencing preparation

Nucleic acid extraction was performed with a lysis buffer containing 2% CTAB (Cetyltrimethylammonium Bromide), 1M NaCl and 100mM Tris-HCl pH8 and purified with the NucleoSpin clean up kit (Macherey-Nagel). Additionally, for each sample, the viral DNA was amplified using illustra GenomiPhi V2 DNA amplification kit according to the manufacturer’s instructions, in order to assure that small circular ssDNA viruses were enriched. Sequencing libraries were constructed using MGIEasy FS Library Prep kit (MGI). DNA was sequenced on a MGI G400 sequencer following the 2x200bp PE protocol at the EcogenO platform (OSERen, Rennes).

### 2.4. Quality control of reads, assembly and annotation

Raw short reads were preprocessed using rqcfilter2 from BBmap (Bushnell., 2014) for adapter and quality trimming. Each viral metagenome was assembled separately with metaSPAdes (Nurk *et al*., 2017). For each date and each sample from the same location, replicates were also co-assembled using metaviralSPAdes (Antipov *et al*., 2020), with parameters of up to 80 threads (-t 80) and 1000 Gb Ram (-m 1000).

### 2.5. Microvirus genomes filtering

To be considered as complete microvirus genomes, assembled contigs need to: (i) have a length between 2,000 nt and 10,000 nt, (ii) be circular (>95% identity between a terminal fragment at the 5′ end and a matching fragment at the 3′ end) and (iii) encode proteins homolog to the Major Capsid Protein (Cap) and to the Replication-Associated Protein (Rep) of known microviruses. For the last criterion, contigs were compared, using a BLASTx strategy using MMseqs2 with a bit-score threshold of 50 (Steinegger and Söding, 2017), to the Cap and Rep protein sequences of 115 microvirus reference genomes (Black *et al*., 2026). These contigs were then deduplicated using a threshold of 100% identity on more than 99.9% reciprocal coverage computed from BLASTn comparisons.

### 2.6. Microvirus genome clustering

Microvirus genomes were grouped into units at various taxonomic levels using their intergenomic similarities computed from BLASTn comparisons as in VIRIDIC (Moraru *et al*., 2020). First, genomes were grouped into genomovars, *i.e.* intra-species units (Aldeguer-Riquelme *et al*. 2024), considering whole-genome average nucleotide identity (ANI) values greater than 99.5% alongside more than 99% of pairwise aligned genomes. Next, genomes were clustered using MCL with the inflation parameter set to 2 (Enright *et al*. 2002) into genus or species level using thresholds recently defined by the ICTV of respectively 50 and 80 on the VIRIDIC intergenomic score. One representative genome was selected for each genus, corresponding to the sequence most closely related to the other members of the group based on the sum of VIRIDIC similarity scores. Representative genomes from all 1,647 genera are available in GenBank under accession number PZ936281-PZ937927.

### 2.7. Microvirus phylogeny and taxonomic assignment

For representative genomes, protein-coding genes were predicted using Prodigal (-p meta: Hyatt *et al*., 2010). Major capsid proteins were identified using BLASTp-like comparisons, performed by MMseqs, between all predicted proteins and Cap proteins from the 115 microviral genera currently defined by the ICTV (Black *et al*. 2026). Cap protein sequences of the current dataset, added to 115 Cap protein sequences of references genomes, were aligned using Clustal Omega using five iterations (–iter=5) (Sievers *et al*., 2011). The alignment was trimmed with TrimAl using a gap threshold of 0.2 (-gapthreshold 0.2) (Capella-Gutiérrez *et al*., 2009), resulting in a trimmed alignment of 638 retained positions. The phylogenetic tree was then inferred using IQ-TREE2 (Minh *et al*., 2020), with the Variable Time model -st AA -m VT+F -bb 1000 -alrt 1000 (Dos Santos *et al*., 2025). The tree was visualized with iTol tool (Letunic and Bork, 2024). The taxonomic affiliation of the genomes at the family and order levels was determined based on the identification of monophyletic groups and the presence of references genomes within those groups.

### 2.8. Relative abundance of microviruses and ecological statistics

All virome reads were mapped onto all complete microvirus genomes with BWA-MEM2 (Vasimuddin *et al*., 2019). The relative abundance of genomes was estimated using CoverM (Aroney *et al*., 2025). In order to assure robust detection, only genomes covered by at least 50% of their length in reads from a given sample were considered present in that sample. Subsequently, genomes were grouped by taxonomic association at the genus level, and their abundances were aggregated. Counts were normalized within each sample (genus counts divided by the total counts of a given sample) and further transformed with Centered Log-Ratio (CLR) method, as recommended for the statistical analysis of metagenomic data (Gloor *et al.,* 2017). Then, alpha diversity of microviral communities was assessed using four complementary indices: Shannon diversity, Chao1 richness, Simpson diversity, and Pielou’s evenness. A linear model of the alpha diversity was constructed for each index with site and season as variables. Normality of residuals was tested using Shapiro test from base R, and homoscedasticity was evaluated using DHARMa package in R. As homoscedasticity was not respected, non-parametric tests were applied. Differences in alpha diversity across seasons and water type were assessed using Kruskal–Wallis tests, followed by pairwise Dunn post-hoc tests with Benjamini–Hochberg correction. Statistical significance was set at p < 0.05.

The beta diversity of the samples was analyzed using CLR data aggregated at the microvirus family level (Gloor *et al*., 2017). The Euclidean distance matrix was calculated using the dist() function from the vegan package. The effect of sampling site and season on microvirus composition was assessed using PERMANOVA via adonis2() (vegan, permutations = 999). The homogeneity of variances across groups was tested using betadisper() and anova() to ensure the validity of the PERMANOVA. Additionally, an analysis of similarities (ANOSIM) was performed to further assess differences in community structure between sites. Pairwise tests between water type were performed using pairwise.adonis2(), and the p-values adjusted according to the Benjamini-Hochberg method were plotted as a heatmap. The overall structure of the samples was explored using principal component analysis (PCA) with prcomp(), and the scores were merged with the metadata for visualization (ggplot2). Confidence ellipses were then drawn to illustrate sample dispersion within each water type, using a 68% confidence interval. The families contributing most to PC1 were identified based on the loadings (pca$rotation). Finally, the variation in PC1 across locations was represented by a boxplot using geom_boxplot() and geom_jitter(), and significant differences were tested using stat_compare_means() (ggpubr, Kruskal-Wallis test). All statistical analyses were performed using R (version 4.3.1).

The 50 most abundant genomes in each of the three specific environments (SW, TW, RW) were selected, resulting in a subgroup of 102 microvirus genomes. A new phylogenetic tree was constructed based on 615 positions retained after trimming following the parameters described above. ORF prediction was processed using prodigal and sequences were annotated using Gaia (Jha *et al*., 2025). Only annotation with high embedding similarity (>90%) and significant alignment (>20% identity and >20% coverage) were conserved.

### 2.9. Virus-Host association

Host associations were inferred using SpaceExtractor (Roux *et al*., 2025) on the 18,449 complete microvirus genomes. In this tool, CRISPR repeats were first identified in assembled public metagenomes and were then used as seeds to identify spacers in metagenomic reads. After a comparison between microvirus genomes and all spacers of the database, a microvirus–repeat pair was considered valid when at least one of the following criteria was met: (i) the microvirus genome has similarity to at least 10 spacers associated to the same repeat, with 0 or 1 mismatch; or (ii) the microvirus genome has similarity to at least 5 spacers with 0–1 mismatch, 10 spacers with 2 mismatches and 20 spacers with 3 mismatches from the same repeat. Bacterial phylogeny was performed using the ape R package by pruning the GTDB tree reference with keep.tip() (bac120_r220.tree ; Parks *et al*., 2022), conserving only the matched hosts.

## 3. RESULTS

### 3.1. Identification of microvirus genomes and fine-scale taxonomic classification

A total of 11.96 billion high-quality reads were generated, with an average length of 198 bp. Independent and cross-assembly of these viromes yielded 76,772 circular contigs ranging in length from 2,000 to 10,000 nt. Among these contigs, 25,533 microvirus genomes were identified by the presence of homologs of both the Major Capsid Protein (Cap) and the Replication-Associated Protein (Rep). Following sequence deduplication, the remaining 19,271 distinct complete genomes were grouped into 15,391 genomovars. These genomovars were clustered into higher taxonomic ranks, resulting in 2,990 species-level vOTU and 1,647 genera using threshold of 80 and 50, respectively, on the intergenomic VIRIDIC (Moraru *et al*., 2020). In order to analyze recurrent subgroups, only genera with at least two genomovars were retained for further analysis, representing 18,449 distinct genomes, 14,569 genomovars, 2,168 species-level vOTU and 825 genera.

### 3.2. Microvirus diversity across the sewage water - treated water - river water continuum

#### 3.2.1. Taxonomic diversity of microvirus

In accordance with the ICTV recommendations, the taxonomic classification of microviruses was based on an analysis of their capsid protein. The phylogeny of the major capsid protein, constructed using one representative sequence from each of the 825 identified genera together with 115 reference sequences, enabled the taxonomic classification of 18,449 genomes (Figure S1). At the highest taxonomic level, genomes are affiliated to the seven defined orders of the class *Microviricetes* (Figure 2). Most of the genomes are affiliated to the orders *Gokushovirales* (15,841 genomes), followed by *Alpavirales* (1,211 genomes), *Secretvirales* (678 genomes), *Amoyvirales* (454 genomes), *Bullavirales* (208 genomes), *Reekeekeevirales* (43 genomes) and *Roodoodoovirales* (14 genomes) (Figure S1).

**Figure 2.**
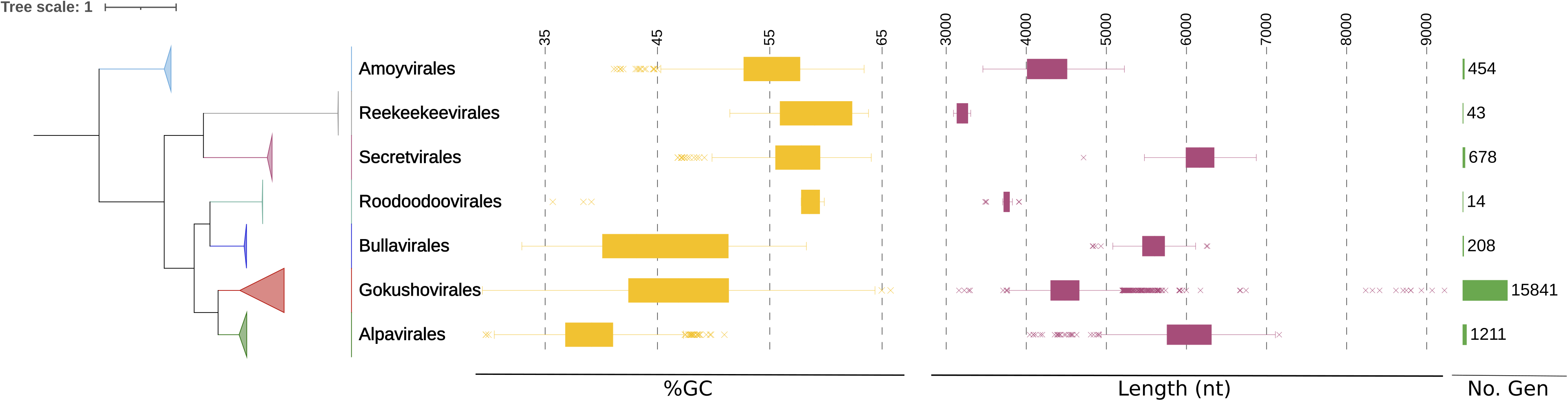
Major capsid protein (Cap) phylogeny and characteristics of microvirus genomes for the seven orders. The affiliation of the SW-TW-RW continuum derived genomes was based on major capsid protein phylogeny (Maximum likelihood, 638 retained positions after trimming, 1000 iterations, VT model). The 825 representative sequences plus 115 compiled reference sequences are included in the tree. On the right are indicated the average GC content (% GC), the average genome length (nt) and the number of newly identified genomes within each order (No. Gen).

While the GC content and the genome length varied strongly within *Microviricetes* orders, some distinct patterns in GC content were observed, with *Alpavirales* exhibiting the lowest average GC content (39.33%) and *Reekeekeevirales* the highest (58.65%) (Figure 2). Average genome lengths ranged from 3,203 nucleotides (nt) in *Reekeekeevirales* to 6,166 nt in *Secretvirales*. These results highlight the broad diversity of microviruses and allow to access their genomic characteristics across the SW-TW-RW continuum of the Seine River.

#### 3.2.2. Variation in microvirus alpha diversity across water types

In order to assess the influence of seasonal variations or water type along the SW-TW-RW continuum, we evaluated the alpha diversity of microvirus communities using several complementary diversity indices (Figure 3 and Figure S2). First, the Chao1 index, showed significant variation between seasons (KW, p = 0.035). Dunn’s post hoc analyses showed significant differences between March and September, with highest richness in March compared to September (p < 0.05), suggesting seasonal shifts in species richness. No significant seasonal effect was detected with Shannon (Kruskal–Wallis test, p=0.22), Simpson (p=0.13) and Pielou (p=0.20) indices.

**Figure 3.**
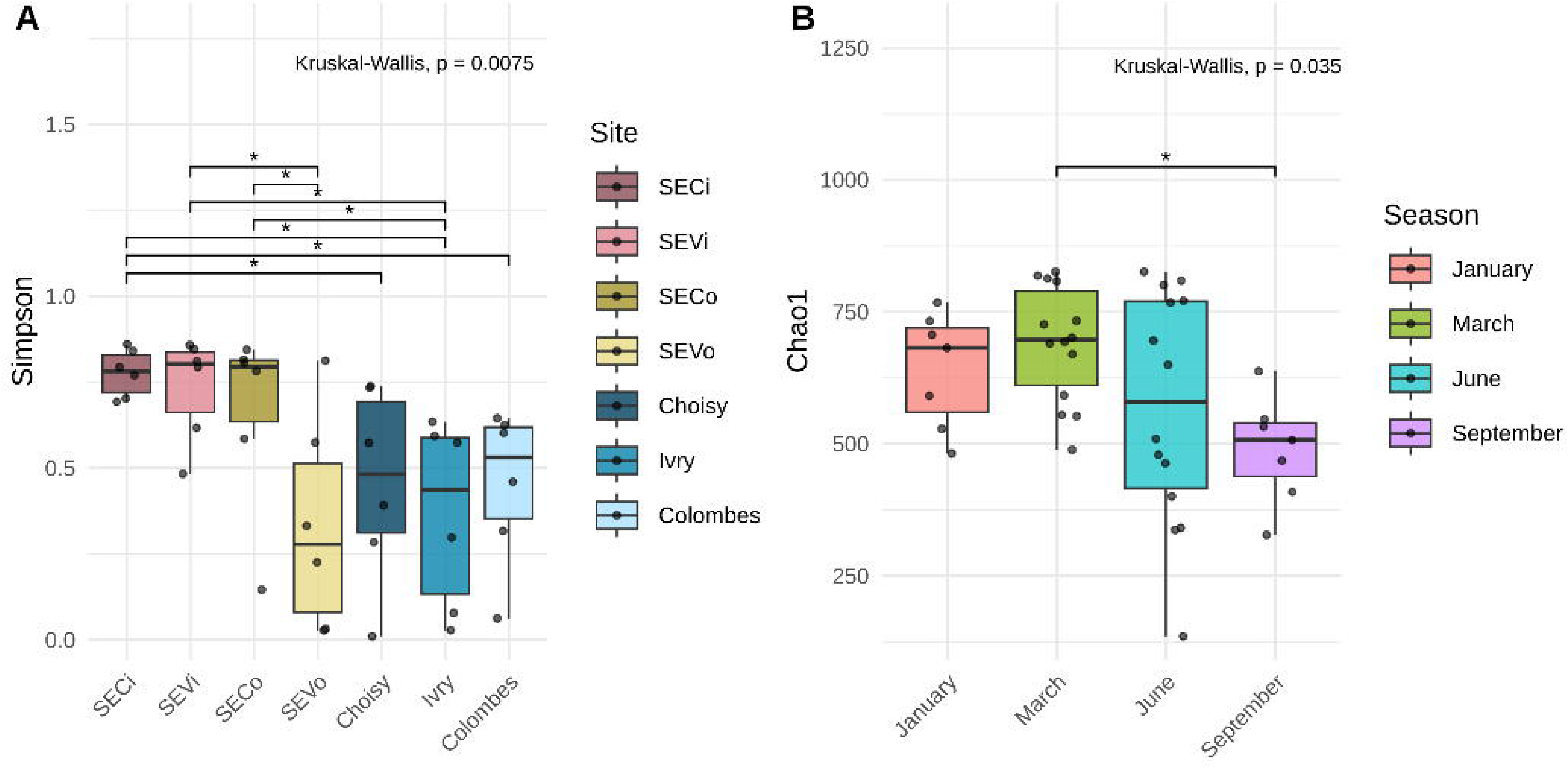
Alpha diversity of the microviral community. Simpson index was estimated according to site (A) and Chao1 according to season (B). Kruskal-Wallis test was performed and significant differences are indicated by stars (p<0.05).

With regard to water type, alpha diversity showed significant differences according to the Shannon (p = 0.0077), Simpson (p = 0.0075) and Pielou (p = 0.0078) indices, but not according to the Chao1 index (p = 0.69). According to Dunn’s post hoc analyses, all indices were significantly higher in sewage water (SW) than in river water (RW), indicating reduced diversity in the receiving environments. When treated water (TW) is considered as a whole, it does not differ from SW and RW, but this is explained by the large difference between the two treatment plants, SEC and SEV. Whilst the diversity of treated water for SEC remained as high as that of SW, that of treated water for SEV was as low as that of RW according to the Shannon, Simpson and Pielou indices. These results indicate marked spatial heterogeneity within the microviral community.

### 3.3. Microvirus community drivers

To investigate patterns in microvirus community structure across the SW–TW–RW continuum, the microviral beta diversity was assessed at the taxonomic family level (Figure 4). Using Principal Component Analysis (PCA), the two main axes deduced explained nearly 80% of the variation of microviral family composition at the different sites, the main axes, PC1, representing 70.2% (Figure 4A). Statistical analyses supported the observed patterns in microvirus community composition across the SW–TW–RW continuum. PERMANOVA revealed significant differences in beta diversity among the seven individual sampling sites (p < 0.001), indicating a strong spatial structuring of viral communities. Pairwise comparisons (Figure 4B) showed no significant differences observed among the three RW indicating a high degree of similarity among the river water samples. This suggests a relative homogeneity of microvirus communities within river environment. In contrast, both SW and TW samples showed distinct community compositions compared with most other sites, including samples belonging to the same water category. The betadisper test showed no significant difference in dispersion between sites (p = 0.53), indicating that the observed significance is not due to unequal variability between groups. ANOSIM confirmed this separation (R = 0.47, p = 0.001), indicating that communities within the same site are more similar to each other than between different sites. Kruskall-Wallis test (p<0.001) on PC1 variation illustrates the marked separation of samples by their water type (Figure 4C). Microvirus families influencing PC1 variance were identified by correlation (Figure 4D). Correlations with PC1 ranged from -0.18 to +0.45, indicating that some microvirus were slightly associated with sites characterized by negative PC1 scores, while others contributed positively. The most pronounced negative correlations were observed for *Pequenoviridae* (-0.18), *Osikeviridae* (-0.14), and *Betagokushoviridae* (-0.12), while the strongest positive contributions came from *Pieniviridae* (+0.45), *Lilleviridae* (+0.43), and *Litenviridae* (+0.33). In general, the coefficients remained modest, suggesting that the effect of each individual viral family on site segregation according to PC1 is weak to moderate, but that the combination of these variations contributes to structuring the beta-diversity observed between water type. Overall, beta diversity analysis indicates that microvirus composition differs significantly by water type, with a clear community structure reflected by PC1 and pairwise comparisons, while intra-site variability remains homogeneous.

**Figure 4.**
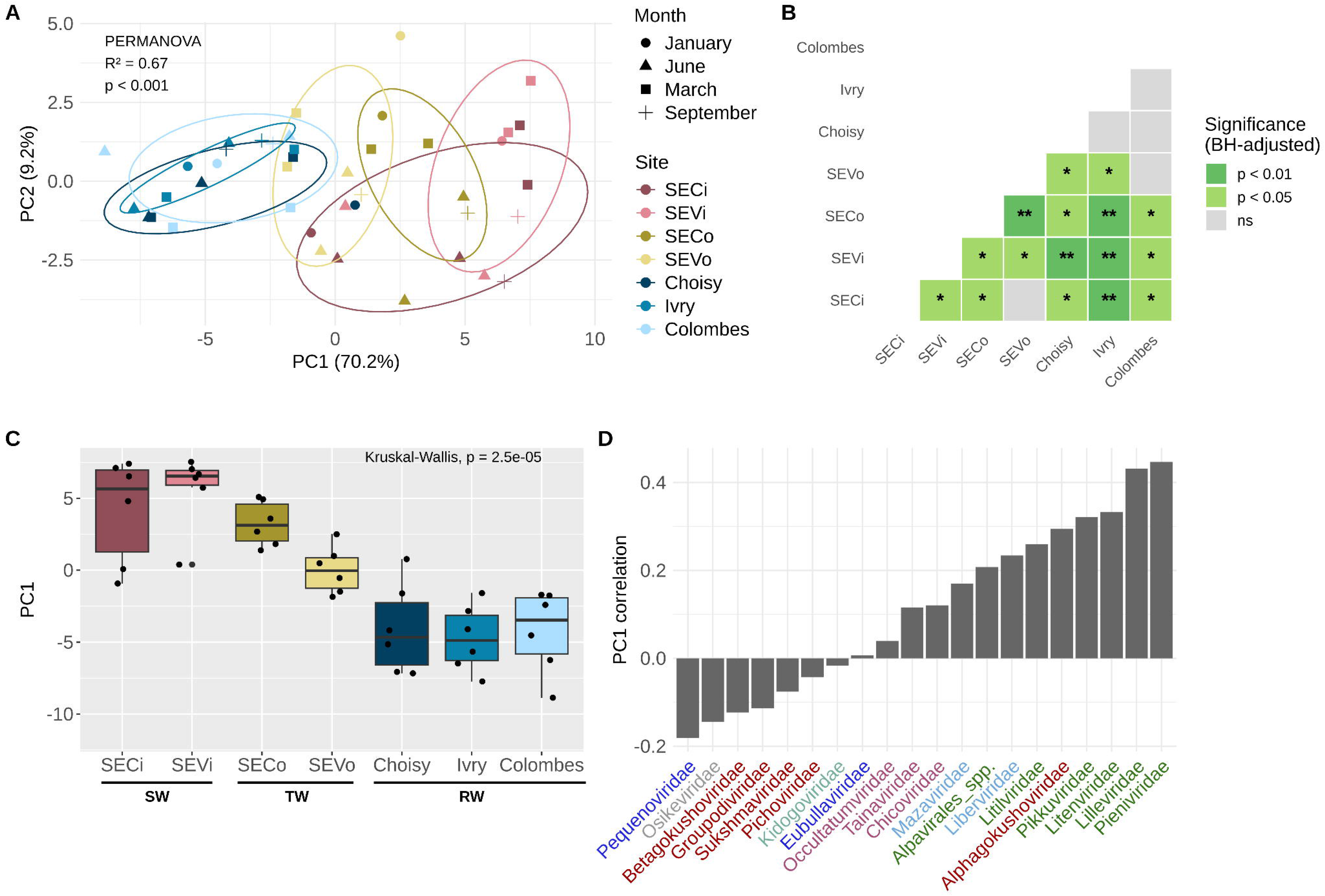
Beta diversity of the microviral community. Analysis was assessed by (A) Principal Component Analysis (PCA), based on Centered-Log Ratio transformed abundances. Each point represents a sample colored according to sampling site and shaped according to sampling month. Ellipses represents confidence intervals at 68% for each site. (B) Heatmap of p-value of pairwise test on PERMANOVA, stars indicates threshold significativity (*p<0.05 , **p<0.01). (C) PC1 scores distribution according to sites. Global differences were tested with Kruskal-Wallis (p<0.001). (D) Contribution of microvirus families, influencing PC1 axis. Family names are colored according to their order as previously described in Figure 2 (red for *Gokushovirales*, green for *Alpavirales*, light blue for *Amoyvirales*, dark blue for *Bullavirales*, purple for *Secretvirales*, grey for *Reekeekeevirales* and turquoise for *Roodoodoovirales*).

### 3.4. Microvirus structuration in the sewage-treated water-river continuum

In order to analyze the flux of microvirus families and get precise relative abundances, reads were mapped on complete genomes (19,271) and analyzed in the SW-TW-RW continuum at the family level. The *Betagokushoviridae* family was dominant in RW, representing more than 92% of the community in all three RW samples, and in TW, where it accounted for 94.22% and 78.14% for SEV (SEVo) and SEC (SECo), respectively (Figure 5A and B). In contrast, while also frequent in SW, *Betagokushoviridae* represented a smaller proportion of the community, at 59.03% and 28.95% for SEV (SEVi) and SEC (SECi), respectively. In these SW samples, the second most abundant taxon was the order *Alpavirales*, representing 26% and 37% of the community in SEV and SEC, respectively. WWTP treatment reduced their relative abundance in TW to about 2.88%, and even more so in river water (1.69% on average), suggesting both a treatment effect within the WWTP and a dilution effect of treated water once released into the natural environment. At the family level, *Litenviridae* represented 18.66% and 24.62% of the community in SW from SEV and SEC, respectively. In SEC (TW) and Colombes (RW), its proportion diminished to 2.76% and 1.67%, respectively. In RW, the relative abundance of the other microvirus families, beside the dominant *Betagokushoviridae*, increased slightly from upstream to downstream of Paris (from 3.7% to 7.6%), with no significant change in their relative proportions to one another (Figure 5C). These distribution patterns indicate that the microvirus community composition is strongly water type specific.

**Figure 5.**
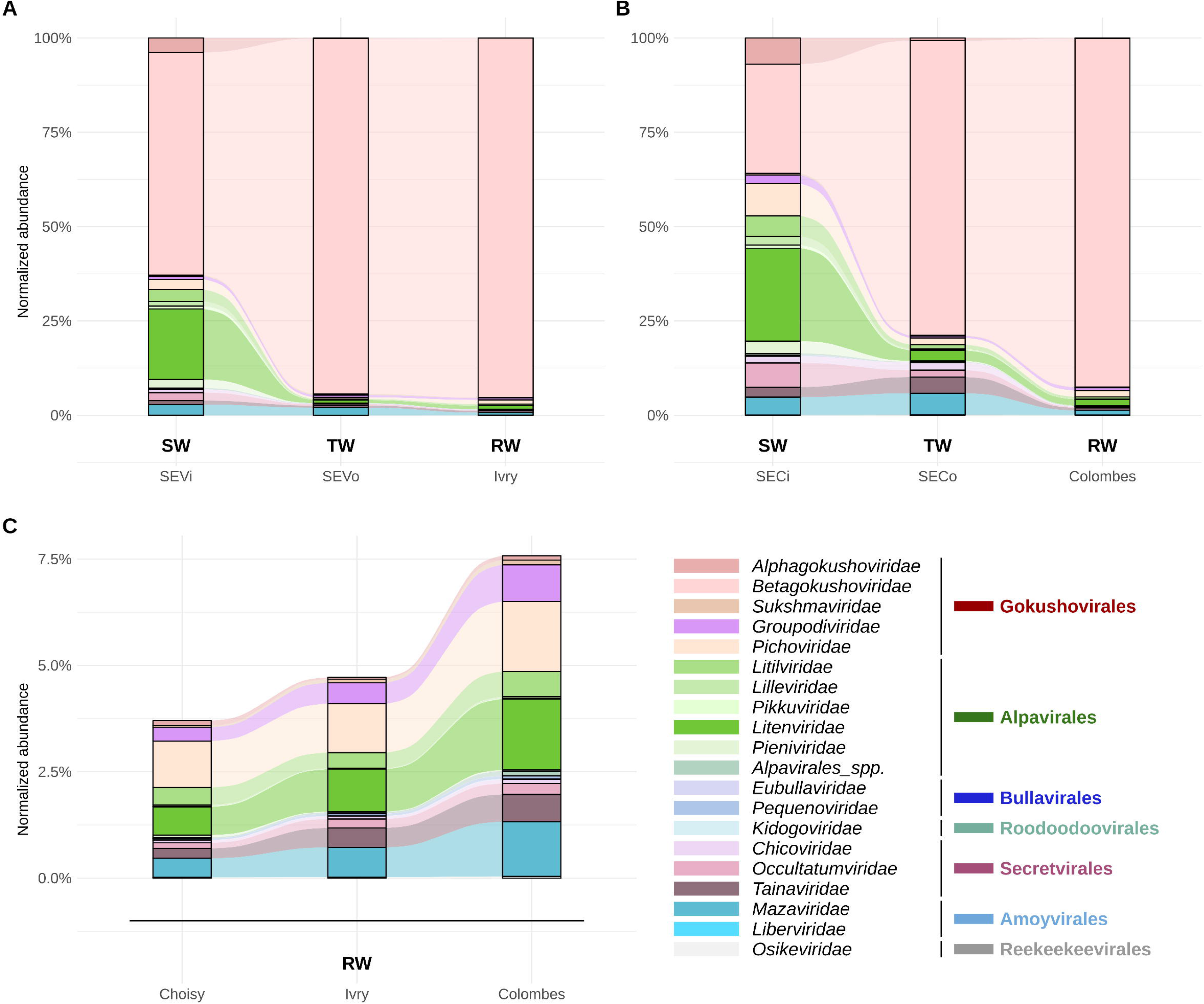
Relative flux of microvirus families across the two SW-TW-RW continuums. along the study area (A and B). (C) shows the relative abundance of these families in RW, focused on non-*Betagokushoviridae*, from upstream to downstream of Paris.

The relative abundance analysis showed distinct signatures depending on the site (Figure 5). In both SW-TW-RW continuums, the SW showed similar community patterns, contrasting with the TW and RW (Figure 5A and 5B). The different community structure observed in TW is associated with the increased dominance of a few persistent viral families, as reflected by their higher proportional abundance.

### 3.5. Most abundant microviruses show distinct relative abundance, genomic and host association patterns

To characterize the diversity and genome structure as well as to assess the persistence throughout the wastewater treatment process, the most abundant microvirus genomes across the entire SW-TW-RW continuum in detail. A subset of 102 microvirus complete genomes, comprising the 50 most abundant genomes in each of the three types of environments (SW, TW, RW), was compiled. Using the major capsid protein phylogeny, these genomes were affiliated to 14 known microvirus families. Five families were affiliated to the *Gokushovirales* order, four to the *Alpavirales*, one to the *Bullavirales*, three to the *Secretvirales*, and one to the *Amoyvirales* (Figure 6). All genomes have a minimum of three protein encoding genes. In addition to the two core genes, namely the Cap and the Rep encoding genes, genomes encoded either a Structural Protein (SP) or a DNA Pilot Protein (DPP) encoding genes. Concerning the structure of the genomes, the arrangement of these three genes varies according to their family. In most genomes, the arrangement is Cap-DPP/SP-Rep. However, genomes assigned to the orders *Secretvirales* and *Amoyvirales*, or to the family *Pichoviridae* within the *Gokushovirales*, exhibit the order Cap-Rep-DPP/SP. DNA Binding Protein and Internal Scafolding Protein encoding genes are only found in the genomes of *Gokushovirales* as already observed previously (Kirchberger and Ochman, 2023), whereas External Scafolding Protein is encoded in *Bullavirales*.

**Figure 6.**
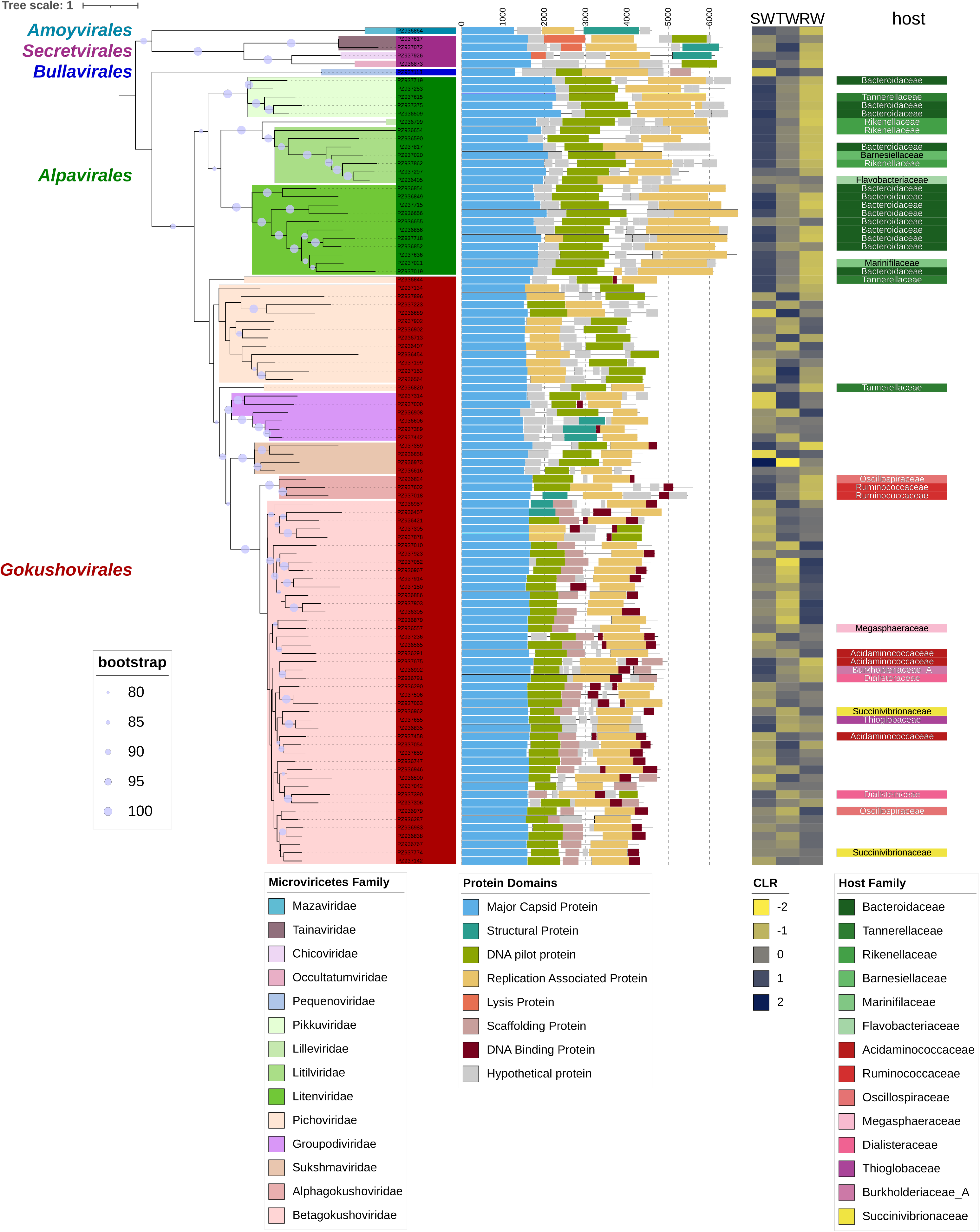
Diversity, relative abundance and host association of most abundant microvirus species in the SW-TW-RW continuum. Phylogenetic tree based on the Cap of 102 most abundant microvirus genomes (Maximum likelihood, 615 retained positions after trimming, VT model). Taxonomy: order level (left), family level (branches). Grey circles indicate bootstrap values with a minimum of 80. The genomic structure of genomes and the annotated protein encoding genes are indicated. The heatmap is based on the CLR transformed abundance and show the difference of relative abundances between SW, TW and RW. Host association at the family level is shown when CRISPR host assignations were robust.

The relative abundance of these microviruses was assessed by the CLR transformed abundance comparing SW, TW and RW. In general, genomes of the four *Alpavirales* families, *Pikkuviridae*, *Lilleviridae*, *Litiviridae* and *Litenviridae,* were most abundant in SW, diminishing in TW and even more in RW (Figure 6). This trend confirms the pattern observed with normalized abundances for the order *Alpavirales* (Figure 5), with the community exhibiting a well-defined structure that differs between sewage water (SW) and river water (RW). For *Gokushovirales* genomes, patterns are mixed with one notable exception at the family level. *Alphagokushoviridae* exhibit the same relative abundance pattern as the *Alpavirales*-affiliated genomes, with high abundance in SW and lower abundance in TW and RW.

### 3.6. Alpavirales and Gokushovirales host association reveals distinct bacterial targets across microvirus orders

Hosts were assigned to microvirus families to identify their host range and to assess host specificity across taxonomic ranks. This approach enables to determine which viruses are associated with single versus multiple hosts and explore patterns of host multiplicity from the species to the phylum level.

Host prediction was assessed by CRISPR spacer matches. From the total of 102 most abundant viral genomes, 43 were linked to at least one potential host. Most genomes affiliated to *Alpavirales* families, particularly present in SW, were predicted to infect hosts belonging to the Bacteroidota phylum (Figure 6), one of the most abundant groups in the human gut (Mahowald *et al*., 2009). In the same vein, genomes belonging to the *Alphagokushoviridae* family, abundant in SW, were predicted to infect different families of the *Eubacteriales* order (Clostridia class and Bacillota phylum), one of the host groups often abundant in the human gut (Salazar-Jaramillo *et al*., 2024). Considering all of the 825 microviral genera, 176 (22%) had at least one predicted host using the CRISPR approach. Of these, 75 microvirus genera were associated with a single host species, 54 were associated with multiple host species and 47 had no species-level attribution for their host. Hosts have been identified only for *Microviricetes* genomes belonging to the two most abundant and diverse orders, the *Alpavirales* and the *Gokushovirales*, and these have revealed specific and different ranges of hosts (Figure S3). Members of the *Gokushovirales* order, the most abundant microviruses in RW and TW, were predicted to infect a wide range of bacterial phyla, indicating substantial host diversity within this group. This pattern is consistent with previous predictions (Labonté *et al*., 2015) and from lytic isolates in *Chlamydia*, *Spiroplasma* and *Bdellovibrio* and temperate in *Enterobacteriaceae* (Kirchberger and Ochman, 2020), which together illustrate that broad host range is a recurring feature among cultured members of this order. Indeed, members of the *Betagokushoviridae* are predicted to infect *Negativicutes* and *Gammaproteobacteria* class belonging to the *Pseudomonadota* phylum, whereas genomes of the *Alphagokushoviridae* family infect bacteria of the *Bacillota* phylum. Interestingly, in relative abundance analysis *Alphagokushoviridae* showed similar patterns as genomes affiliated to *Alpavirales* (Figure 6). *Pichoviridae* genomes was the only family of this order to infect *Bacteroidota* and *Fibrobacterota* phylum and not the *Pseudomonadota* or *Bacillota* phylum.

In contrast, *Alpavirales* genomes were restricted to a single bacterial phylum, predominantly associated with gut-related bacteria, consistent with the higher abundance of this order in SW, and extending the pattern already observed among the 102 most abundant genomes (see above). Members of the *Alpavirales* order mainly infect bacteria of the family *Bacteroidaceae*, consistent with proviruses belonging to this microvirus order (Krupovic and Forterre 2011). All the bacterial families infected by this microvirus order were associated to the *Bacteroidota* phylum, especially *Bacteroidales* order. Members of the *Litenviridae*, which appeared to be the characteristic family of SW, were found to infect three main host genera: *Prevotella*, *Bacteroides* and *Phocaeicola*.

## 4. DISCUSSION

### 4.1 Identification of high number of microvirus genomes in SW-TW-RW continuum allow high taxonomic resolution

Microvirus spatiotemporal dynamics across the continuum from sewage water (SW) to treated wastewater (TW) and river water (RW) remain largely unexplored, particularly at fine taxonomic scales. This study represents a substantial and unprecedented expansion of the complete microvirus genome catalog. Our dataset comprises 19,271 complete microvirus genomes recovered directly from *in situ* environmental samples. Previously, Kircherberger *et al*. (2022) curated a set of 4,077 complete microviral genomes from isolates, metagenome-assembled genomes and prophage sequences, predominantly originating from human, primate, mammalian and marine viromes; a further ∼9,000 genomes were subsequently and non-manually assigned to this taxonomy from additional metagenomic datasets, including 3,871 sequences from a global wastewater dataset (Kirstahler *et* al., 2021). However, freshwater and wastewater treatment plant environments remained essentially underrepresented among their manually curated genomes. By grouping our microvirus genomes into 825 genera, combined with the recently reorganized taxonomy of the microviruses (Black *et al*., 2026), our study provides the first high-resolution, curated view of microviral diversity across an urban aquatic continuum, substantially expanding the taxonomic and ecological representation of these environments.

### 4.2 Microviruses show high taxonomic and genomic diversity across the SW-TW-RW continuum

Previous studies have identified microvirus in marine environments, freshwater systems, peatlands, sediments, animal-associated microbiomes, and wastewater ecosystems (Roux *et al*., 2012; Labonté and Suttle, 2013; Quaiser *et al*., 2015, Kirchberger *et al*., 2022) allowing the delineation in seven microviral orders (Black *et al*., 2026). In this study, we detected microvirus genomes spanning these seven orders and 20 families, within a single urban aquatic continuum, highlighting both, the ecological success and broad environmental distribution of these viruses. This consistency in taxonomic composition suggests a global distribution of major microvirus orders across the SW-TW-RW continuum. The overwhelming dominance of *Gokushovirales* order is consistent with numerous metagenomic surveys. Members of this order are among the most frequently detected microvirus in aquatic environments and have been reported from oceans, lakes, rivers, peatland, groundwater, and wastewater systems (Roux *et al*., 2012; Quaiser *et al*., 2015, Kirchberger and Ochman, 2023). Due to the high number of reconstructed complete microvirus genomes from *in situ* samples, fine scale taxonomic resolution analysis as well as their spatio-temporal dynamic across the SW-TW-RW continuum could be performed. In total, the genomes were affiliated to 20 different microvirus families suggesting a broad host spectrum that contributes to their widespread environmental distribution. Therefore, ecological variations became evident and distinct viral assemblages could be associated with each compartment. Within the *Gokushovirales* order, the *Betagokushoviridae* family represented the dominant family across TW and RW samples. Unlike *Alpavirales* microviruses, whose hosts appear largely restricted to gut-associated bacteria, *Betagokushoviridae* microviruses were predicted to infect bacterial groups commonly found in both wastewater and aquatic environments. Such host flexibility likely contributes to their persistence throughout the continuum.

The genome organization observed in our recovered genomes was consistent with previous observations (Roux *et al*., 2012; Quaiser *et al*., 2015; Doore and Fane, 2016; Kirchberger and Ochman, 2023). All genomes encode a major capsid protein (Cap), a replication initiation protein (Rep), and additional structural proteins. Gene arrangements were largely conserved within families and were consistent with the Cap-based phylogeny (Figure 6). Within the *Gokushovirales* order, the *Betagokushoviridae* family show a specific organization with the presence of a DNA binding protein and a Scaffolding protein (external or internal) in almost all the most abundant genomes. This conservation of genome architecture across environmental compartments suggests that major evolutionary constraints shape microvirus genome organization despite considerable ecological diversification. The broad differences in GC content observed among orders may further reflect long-term adaptation to distinct bacterial hosts: *Gokushovirales* genomes, which infect phylogenetically diverse bacteria, spanned a wide range of GC content, whereas *Alpavirales* genomes, restricted to a single bacterial phylum, showed a comparatively narrower GC range. Previous studies have shown that viral nucleotide composition frequently mirrors host genomic characteristics due to mutational pressures and host-driven selection (Ahlgren *et al.,* 2017). Consequently, the genomic diversity observed here likely reflects the broad diversity of bacterial hosts present along the SW–TW–RW continuum.

### 4.3 Water type outweighs seasonal variation in structuring microvirus communities

One of the major findings of this study is the limited influence of seasonality compared with the strong effect of water type. Despite the exclusive use of complete genomes assembled from *in situ* samples, richness (Chao1) differed only between two sampling periods (March and September) and no significant seasonal differences were detected for Shannon, Simpson, or Pielou indices (Figure S2). Similarly, beta-diversity analyses did not reveal a clear seasonal clustering of samples. At the family level, the model only including month did not show a significant result. At the genus level, the model was significant (Table S1), but after correction, no pairwise comparison remained significant, suggesting that seasonality is a non-robust signal in this dataset (Table S2). However, microvirus ecology is primarily governed by bacterial host dynamics, and the unusually high river discharge observed throughout 2024 likely reduced the detectability of any underlying seasonal signal. Indeed, the year 2024 was characterized by exceptionally high precipitation and elevated Seine River flow rate. Under such conditions, enhanced mixing and homogenization of microbial communities reduce temporal variability by continuously redistributing microorganisms throughout the watershed. High flow conditions may therefore mask seasonal effects that could become apparent during periods of lower discharge. This extensive genome catalog nonetheless provides a valuable resource for reassessing seasonal patterns in future studies or retrospective datasets collected under more typical hydrological conditions.

In contrast, water type emerged as the dominant factor structuring viral communities. Both alpha- and beta-diversity analyses consistently separated SW, TW, and RW samples. This pattern suggests that each compartment represents a distinct ecological niche characterized by different physicochemical conditions, bacterial communities, and viral selection pressures. Interestingly, heterogeneity was observed between the two WWTPs input and output. While SW are expected to reflect similar human-associated microbial sources, the observed differences between SEC and SEV indicate that local catchment characteristics may influence incoming viral communities. Variability in population density, industrial contributions, sewer residence time, or stormwater inputs may contribute to this heterogeneity. The contrast between both of the TW were even more pronounced. These observations likely reflect differences in treatment processes between the two WWTPs. Such results suggest that WWTPs filtering does not lead to a universal TW virome but instead generate site-specific viral communities. Conversely, RW appeared remarkably homogeneous despite their spatial separation along the Seine River. The absence of a clear upstream–downstream trend suggests that viral communities are rapidly homogenized within the river system. This result may reflect the large volume of the Seine River, intense hydrological mixing, and the cumulative influence of multiple anthropogenic inputs. In addition, this could suggest that microviruses infect essentially the most stable, abundant, and prevalent bacterial taxa.

### 4.4 WWTPs act as ecological filters reshaping microvirus communities

The transition from SW to TW represents the strongest ecological shift observed in this study. The decrease of *Alpavirales* throughout treatment illustrates this filtering effect. Members of this order were highly abundant in SW but strongly reduced in TW and RW. CRISPR-based host predictions indicate that these viruses primarily infect *Bacteroidales*, including *Prevotella*, *Bacteroides*, and *Phocaeicola*, which are characteristic members of the human gut microbiota. Several mechanisms may explain this decline. First, wastewater treatment substantially reduces the abundance of strict anaerobic gut-associated bacteria. Consequently, phages dependent upon these hosts may also decline. Second, dilution by environmental viruses becomes increasingly important after discharge into rivers. Third, lysogenic lifestyles may contribute to *Alpavirales* ecology. Krupovic and Forterre (2011) demonstrated that several *Alpavirales*-related microviruses exist as prophages within gut-associated bacteria. The disruption of these host populations during treatment could therefore directly affect the maintenance of associated phages. In contrast, *Betagokushoviridae* family members of the *Gokushovirales* order increased substantially after treatment and dominated river communities. This pattern suggests either persistence during treatment or maintenance through environmentally adapted host populations. Consequently, treatment processes may indirectly favor these viruses by maintaining suitable host populations. Importantly, both alpha- and beta-diversity analyses indicate that treatment generates a distinct viral assemblage. This observation suggests that WWTPs constitute unique ecological habitats capable of selecting specific microviral populations. WWTPs combine aeration, sedimentation, biological degradation, and chemical treatments that profoundly modify microbial ecosystems (Hong *et al*., 2024; Jin *et al*., 2025). Similar conclusions have recently been proposed for bacterial and viral communities in wastewater systems (Wang *et al*., 2020; Guajardo-Leiva *et al*., 2025; Niu *et al*., 2026).

### 4.5 Microviruses could serve as potential bioindicators of anthropized aquatic systems

The strong compartment-specific distribution of microvirus suggests that these viruses may represent valuable indicators of anthropogenic influence. *Alpavirales* order members appear to be reliable markers of wastewater-associated microbial communities because of their strong association with gut bacteria and their high abundance in SW. In contrast, *Betagokushoviridae* family may reflect environmental conditions prevailing in TW and anthropized RW. Current monitoring strategies primarily focus on fecal indicator bacteria or enteric viruses (Farkas *et al.,* 2020). However, bacteriophages provide complementary ecological information because they integrate both host distribution and environmental conditions. Recent studies have already proposed specific phages as indicators of wastewater contamination (Ballesté *et al*., 2022; Bichet *et al*., 2024). The strong and reproducible shifts observed along the SW–TW–RW continuum suggest that particular groups of microvirus could contribute to future biomonitoring approaches, such as *Litenviridae* family. Their high abundance, host association, and sensitivity to treatment-associated processes make them promising candidates for tracking anthropogenic gradients in freshwater ecosystems.

## CONCLUSION

This study provides a high-resolution view of microvirus diversity across an urban sewage–wastewater treatment plant–river continuum and shows that water type is the primary driver of community structure, outweighing seasonal variation under the hydrological conditions encountered. The scale-dependent response highlights the importance of high-resolution taxonomic approaches for understanding freshwater virome ecology and identify ecological patterns associated with anthropogenic pressures. By recovering 18,449 complete microvirus genomes and resolving them into 825 genera, we reveal that urban aquatic compartments host highly differentiated microviral assemblages, despite the broad representation of all currently described microvirus orders. In particular, the decrease of *Alpavirales* from sewage water to treated water and river water and the persistence and dominance of *Betagokushoviridae* in treated and river water, indicates that wastewater treatment acts as a strong ecological filter shaping microvirus communities. These patterns also suggest that microviruses may complement traditional microbial indicators for monitoring wastewater impacts in freshwater systems. Although host assignment remains partial and the study was restricted to a single watershed and a limited number of treatment plants, the observed compartment-specific signatures are robust and ecologically meaningful. Future work should extend this approach to multiple catchments and treatment configurations, integrate host-resolved and functional analyses, and test whether selected microvirus groups can serve as reliable bioindicators of human influence in urban waters.

## Author Contributions

**Alisa Langlais:** Conceptualization, Data curation, Investigation, Methodology, Visualization, Writing – original draft, Writing – review & editing; **Celtill Dumont:** Data curation, Methodology, Writing – original draft, Writing – review & editing; **Jérémy Mougins:** Funding acquisition, Resources, Writing review & editing **Sabrina Guérin-Rechdaoui**: Funding acquisition, Resources; **Céline Roose-Amsaleg:** Conceptualization, Formal analysis, Funding acquisition, Investigation, Methodology, Project administration, Resources, Supervision, Validation, Writing original draft, Writing review & editing; **François Enault:** Data curation, Investigation, Methodology, Supervision, Validation, Writing – original draft, Writing – review & editing; **Achim Quaiser:** Conceptualization, Data curation, Funding acquisition, Investigation, Methodology, Resources, Supervision, Validation, Writing – original draft, Writing – review & editing

## ACKNOWLEDGMENT

We acknowledge the GenOuest bioinformatics core facility (https://www.genouest.org) for providing the computing infrastructure. We thank Sophie Michon-Coudouel, Romain Causse- Védrines and Antoine Peigné from the EcogenO core facility of Rennes (OSERen, Biogenouest, ANAEE-Fr) for their technical support. We also thank Erwan Garcia-Gonzalez, Carlyne Lacroix, Robin Richoux, Véronique Brémont and Anne-Sophie Permal (from the Greater Paris Sanitation Authority - SIAAP) for water sampling and their assistance with water pretreatment.

## SUPPLEMENTARY MATERIALS

**Figure S1 Phylogenetic analysis of full-length Cap sequences.** Included are the 825 sequences at the genus level representing the diversity of microvirus found in the sewage-river continuum and 115 reference genomes. The inner circle shows the taxonomic assignment, based on the position of the genus in the tree, the second circle shows the percentage of GC content and the third circle indicates the length of the genomes. Gaps in the circles correspond to the reference genomes.

**Figure S2 Alpha-diversity indices, according to site (left) or season (right).** Shannon (A, B), Simpson (C, D) and Pielou (E, F) and Chao1 (G,H) indices were estimates. Differences among groups were tested with Kruskal–Wallis tests (p-values are indicated in each panel). Brackets with an asterisk indicate pairwise differences identified by Dunn’s post hoc test (Benjamini–Hochberg adjusted p ≤ 0.05). SECi and SEVi: WWTP inlets (sewage water); SECo and SEVo: WWTP outlets (treated water); Choisy, Ivry and Colombes: river water.

**Figure S3 Abundance of CRISPR-based host prediction of *Microviricetes* genomes affiliated to the orders *Gokushovirales* and *Alpavirales*.** The tree represents the bacterial hosts (tree pruned from bac120_r220.tree, GTDB database). Different phyla, classes and orders of bacterial hosts are highlighted. Darker boxes indicate a higher number of validated CRISPR spacer matches between viral genomes and host-associated spacer sequences.

**Table S1** Results of PERMANOVA analyses (PERmutational Multivariate ANalysis of VAriance, based on a Bray-Curtis dissimilarity matrix, 999 permutations) testing the effect of sampling site (Location), sampling month (Month), and their combination (Loc + Month) on community composition, at the family and genus taxonomic levels. RZ: proportion of variance explained by the factor; p_value: associated p-value; Signif: significance threshold (** p < 0.01; * p < 0.05).

**Table S2** Pairwise comparisons (pairwise PERMANOVA, Bray-Curtis dissimilarity, 999 permutations) between sampling site (Location) and between sampling months (Month), for the Location, Month, and Loc + Month factors, at the family and genus taxonomic levels. RZ: proportion of variance explained; p_value: raw p-value; p_adj: p-value adjusted for multiple comparisons (Benjamini-Hochberg method); Signif: significance threshold (** p_adj < 0.01; * p_adj < 0.05; . p_adj < 0.1).

