## Supplementary material for "Unprecedented urban freshwater microviral survey highlights shift from sewage to river water communities": Figure S1

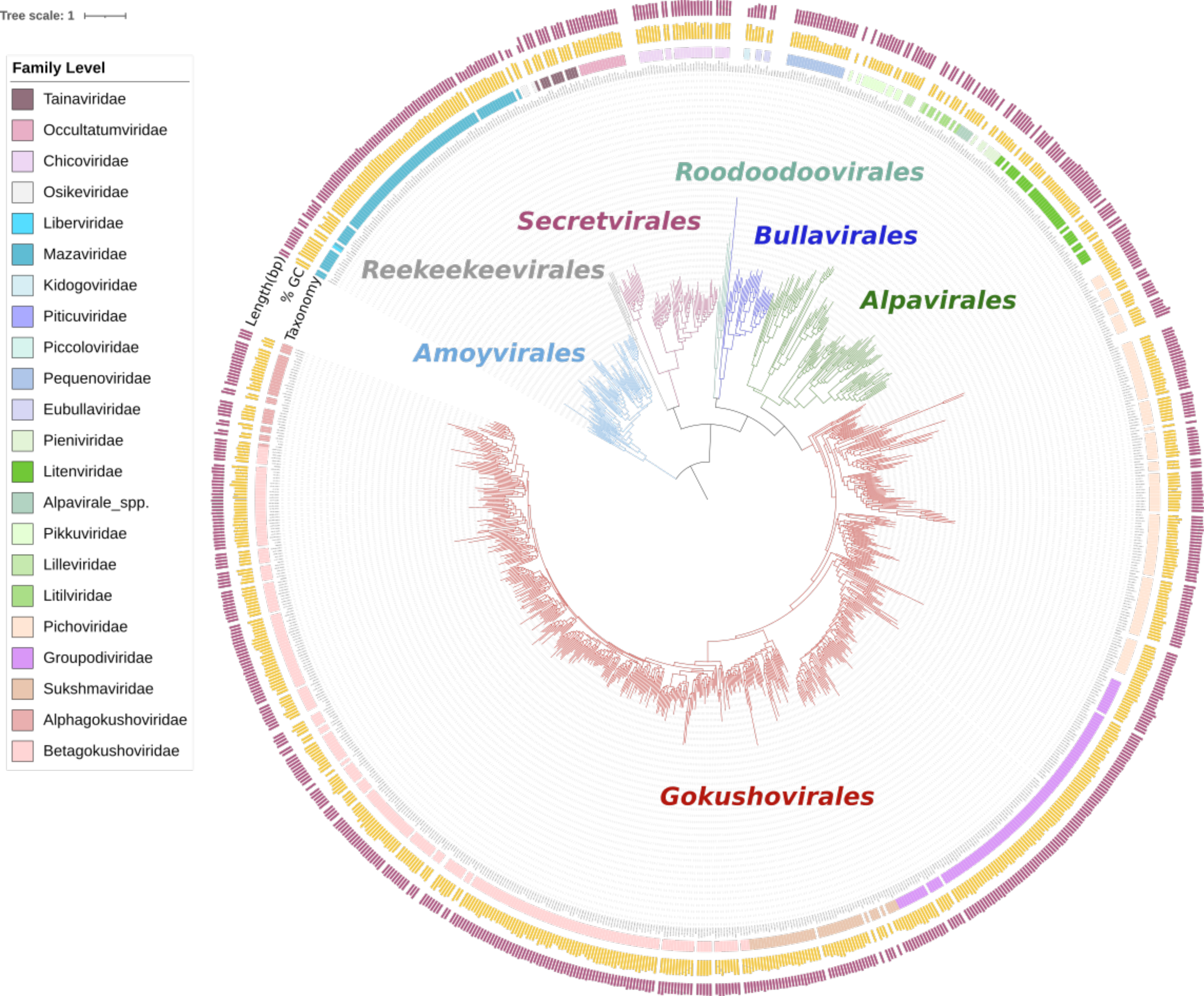

Tree scale: 1
Family Level
Tainaviridae
Occultatumviridae
Chicoviridae
Osikeviridae
Liberviridae
Mazaviridae
Kidogoviridae
Piticuviridae
Piccoloviridae
Pequenoviridae
Eubullaviridae
Pieniviridae
Litenviridae
Alpavirale_spp.
Pikkuviridae
Lilleviridae
Litilviridae
Pichoviridae
Groupodiviridae
Sukshmaviridae
Alphagokushoviridae
Betagokushoviridae
Length(bp)
% GC
Taxonomy
Roodoodoovirales
Secretvirales
Bullavirales
Reekeekeevirales
Alpavirales
Amoyvirales
Gokushovirales
