## Supplementary material for "Unprecedented urban freshwater microviral survey highlights shift from sewage to river water communities": Figure S2

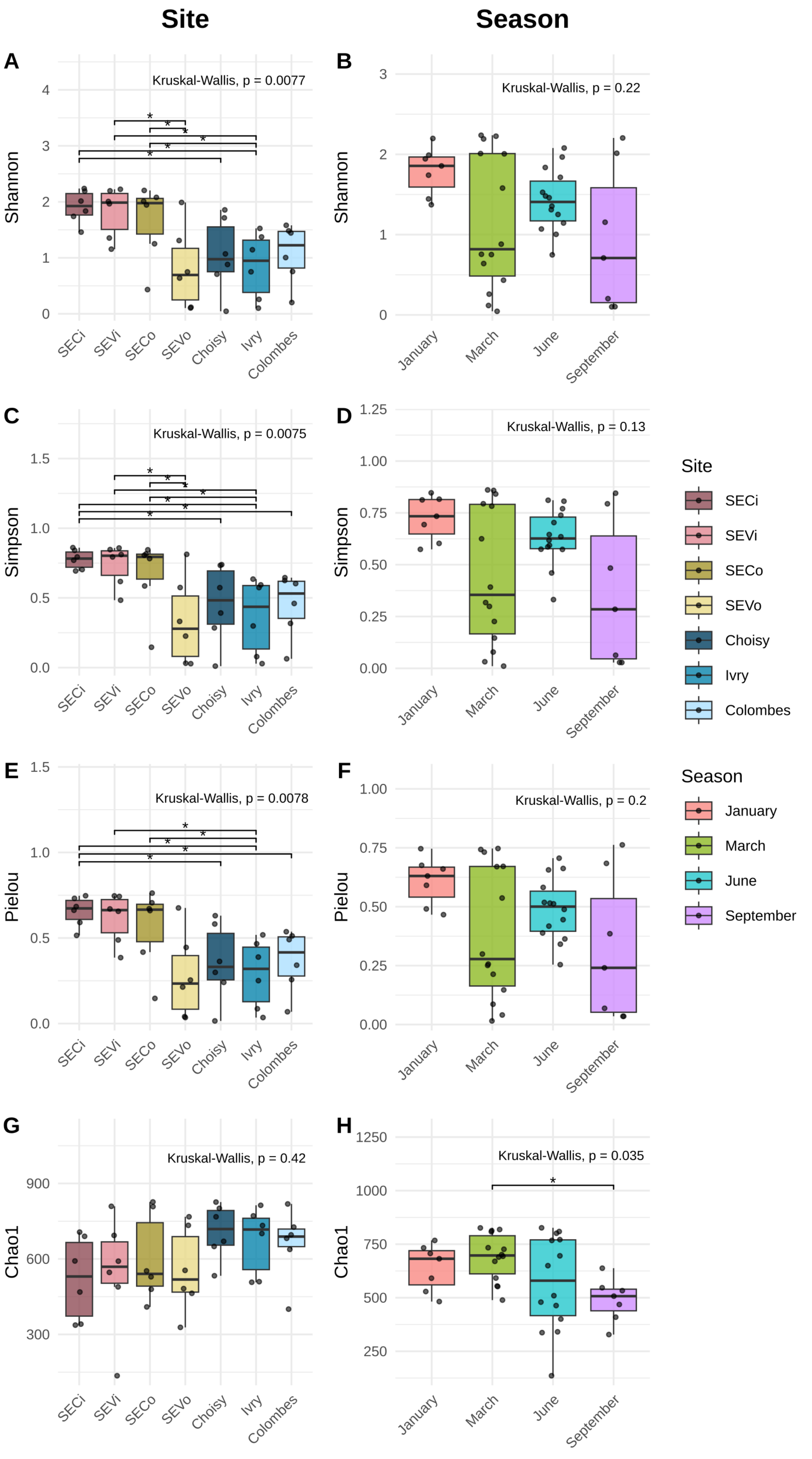

Site
Season
A
Kruskal-Wallis, p = 0.0077
Shannon
SECi
SEVi
SECo
SEVo
Choisy
Ivry
Colombes
B
Kruskal-Wallis, p = 0.22
January
March
June
September
C
Kruskal-Wallis, p = 0.0075
Simpson
D
Kruskal-Wallis, p = 0.13
E
Kruskal-Wallis, p = 0.0078
Pielou
F
Kruskal-Wallis, p = 0.2
G
Kruskal-Wallis, p = 0.42
Chao1
H
Kruskal-Wallis, p = 0.035
Site
SECi
SEVi
SECo
SEVo
Choisy
Ivry
Colombes
Season
January
March
June
September
