## Supplementary material for "Unprecedented urban freshwater microviral survey highlights shift from sewage to river water communities": Figure S3

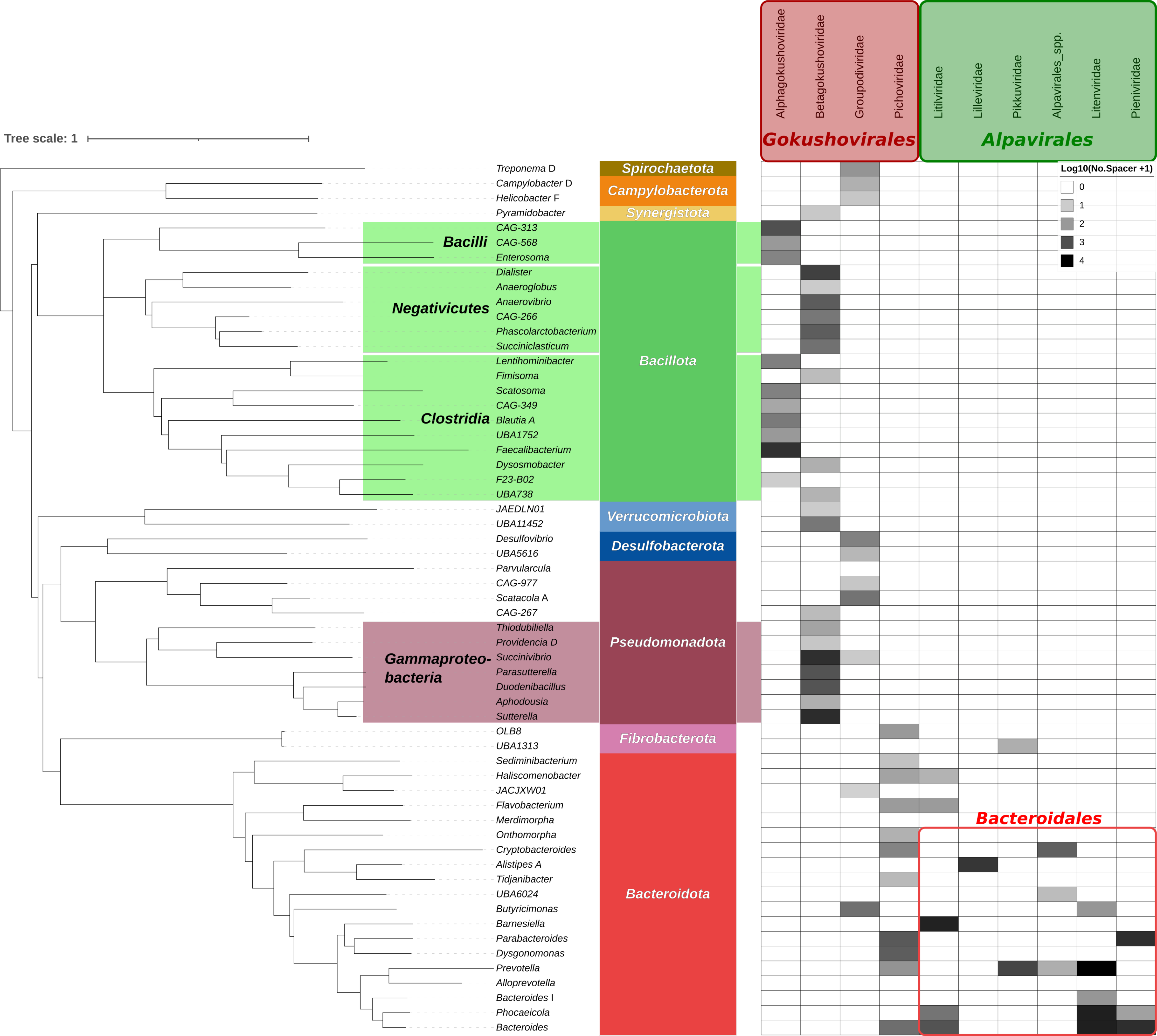

Tree scale: 1
Gokushovirales
Alphagokushoviridae
Betagokushoviridae
Groupodiviridae
Pichoviridae
Alpavirales
Litilviridae
Lilleviridae
Pikkuviridae
Alpavirales_spp.
Litenviridae
Pieniviridae
Log10(No.Spacer +1)
0
1
2
3
4
Spirochaetota
Campylobacterota
Synergistota
Bacillota
Verrucomicrobiota
Desulfobacterota
Pseudomonadota
Fibrobacterota
Bacteroidota
Bacilli
Negativicutes
Clostridia
Gammaproteo-bacteria
Bacteroidales
Treponema D
Campylobacter D
Helicobacter F
Pyramidobacter
CAG-313
CAG-568
Enterosoma
Dialister
Anaeroglobus
Anaerovibrio
CAG-266
Phascolarctobacterium
Succiniclasticum
Lentihominibacter
Fimisoma
Scatosoma
CAG-349
Blautia A
UBA1752
Faecalibacterium
Dysosmobacter
F23-B02
UBA738
JAEDLN01
UBA11452
Desulfovibrio
UBA5616
Parvularcula
CAG-977
Scatacola A
CAG-267
Thiodubiliella
Providencia D
Succinivibrio
Parasutterella
Duodenibacillus
Aphodousia
Sutterella
OLB8
UBA1313
Sediminibacterium
Haliscomenobacter
JACJXW01
Flavobacterium
Merdimorpha
Onthomorpha
Cryptobacteroides
Alistipes A
Tidjanibacter
UBA6024
Butyricimonas
Barnesiella
Parabacteroides
Dysgonomonas
Prevotella
Alloprevotella
Bacteroides I
Phocaeicola
Bacteroides
