## Supplementary material for "Unprecedented urban freshwater microviral survey highlights shift from sewage to river water communities": Table S1

| *Tax level* | *Factor* | *R2* | *p_value* | *Signif* |
|---|---|---|---|---|
| **Family** | **Location** | **0,583** | **0,001** | ** |
| Family | Month | 0,084 | 0,322 | |
| **Family** | **Loc + Month** | **0,667** | **0,001** | ** |
| **Genus** | **Location** | **0,362** | **0,001** | ** |
| **Genus** | **Month** | **0,133** | **0,008** | ** |
| **Genus** | **Loc + Month** | **0,496** | **0,001** | ** |
