## Supplementary material for "Unprecedented urban freshwater microviral survey highlights shift from sewage to river water communities": Table S2

| *Tax level* | *Factor* | *Comparison* | *R2* | *p_value* | *p_adj* | *Signif* |
|---|---|---|---|---|---|---|
| **Family** | **Location** | **SEVo_vs_Ivry** | **0,379** | **0,004** | **0,01** | * |
| **Family** | **Location** | **SEVo_vs_SEVi** | **0,403** | **0,005** | **0,01** | * |
| **Family** | **Location** | **SEVo_vs_SECo** | **0,306** | **0,002** | **0,01** | * |
| **Family** | **Location** | **Choisy_vs_SECi** | **0,508** | **0,005** | **0,01** | * |
| **Family** | **Location** | **Choisy_vs_SEVi** | **0,632** | **0,005** | **0,01** | * |
| **Family** | **Location** | **Choisy_vs_SECo** | **0,545** | **0,006** | **0,01** | * |
| **Family** | **Location** | **Ivry_vs_SECi** | **0,585** | **0,002** | **0,01** | * |
| **Family** | **Location** | **Ivry_vs_SEVi** | **0,695** | **0,004** | **0,01** | * |
| **Family** | **Location** | **Ivry_vs_SECo** | **0,639** | **0,002** | **0,01** | * |
| **Family** | **Location** | **Colombes_vs_SECi** | **0,524** | **0,004** | **0,01** | * |
| **Family** | **Location** | **Colombes_vs_SEVi** | **0,642** | **0,001** | **0,01** | * |
| **Family** | **Location** | **Colombes_vs_SECo** | **0,556** | **0,003** | **0,01** | * |
| **Family** | **Location** | **SEVi_vs_SECo** | **0,269** | **0,006** | **0,01** | * |
| **Family** | **Location** | **SEVo_vs_Colombes** | **0,315** | **0,007** | **0,011** | * |
| **Family** | **Location** | **SEVo_vs_Choisy** | **0,297** | **0,013** | **0,018** | * |
| **Family** | **Location** | **SEVo_vs_SECi** | **0,285** | **0,024** | **0,032** | * |
| Family | Location | SECi_vs_SECo | 0,132 | 0,178 | 0,22 | |
| Family | Location | SECi_vs_SEVi | 0,098 | 0,327 | 0,382 | |
| Family | Location | Choisy_vs_Ivry | 0,047 | 0,743 | 0,821 | |
| Family | Location | Ivry_vs_Colombes | 0,032 | 0,902 | 0,947 | |
| Family | Location | Choisy_vs_Colombes | 0,022 | 0,98 | 0,98 | |
| **Family** | **Loc + Month** | **SEVo_vs_SECo** | **0,649** | **0,001** | **0,007** | ** |
| **Family** | **Loc + Month** | **Choisy_vs_SEVi** | **0,792** | **0,001** | **0,007** | ** |
| **Family** | **Loc + Month** | **Ivry_vs_SECo** | **0,762** | **0,001** | **0,007** | ** |
| **Family** | **Loc + Month** | **Ivry_vs_SECi** | **0,784** | **0,002** | **0,008** | ** |
| **Family** | **Loc + Month** | **Ivry_vs_SEVi** | **0,813** | **0,002** | **0,008** | ** |
| **Family** | **Loc + Month** | **SEVi_vs_SECo** | **0,587** | **0,004** | **0,014** | * |
| **Family** | **Loc + Month** | **SEVo_vs_SEVi** | **0,669** | **0,006** | **0,018** | * |
| **Family** | **Loc + Month** | **SEVo_vs_Choisy** | **0,627** | **0,01** | **0,023** | * |
| **Family** | **Loc + Month** | **Colombes_vs_SEVi** | **0,744** | **0,01** | **0,023** | * |
| **Family** | **Loc + Month** | **Choisy_vs_SECo** | **0,687** | **0,014** | **0,026** | * |
| **Family** | **Loc + Month** | **Colombes_vs_SECi** | **0,696** | **0,015** | **0,026** | * |
| **Family** | **Loc + Month** | **SECi_vs_SECo** | **0,556** | **0,014** | **0,026** | * |
| **Family** | **Loc + Month** | **Colombes_vs_SECo** | **0,662** | **0,018** | **0,029** | * |
| **Family** | **Loc + Month** | **SECi_vs_SEVi** | **0,589** | **0,021** | **0,032** | * |
| **Family** | **Loc + Month** | **SEVo_vs_Ivry** | **0,595** | **0,026** | **0,036** | * |
| **Family** | **Loc + Month** | **Choisy_vs_SECi** | **0,668** | **0,032** | **0,042** | * |
| Family | Loc + Month | SEVo_vs_SECi | 0,539 | 0,05 | 0,062 | . |
| Family | Loc + Month | SEVo_vs_Colombes | 0,497 | 0,12 | 0,14 | |
| Family | Loc + Month | Choisy_vs_Ivry | 0,405 | 0,371 | 0,41 | |
| Family | Loc + Month | Ivry_vs_Colombes | 0,371 | 0,474 | 0,498 | |
| Family | Loc + Month | Choisy_vs_Colombes | 0,339 | 0,555 | 0,555 | |
| Family | Month | January_vs_March | 0,043 | 0,391 | 0,469 | |
| Family | Month | January_vs_June | 0,075 | 0,197 | 0,469 | |
| Family | Month | January_vs_September | 0,078 | 0,318 | 0,469 | |
| Family | Month | March_vs_June | 0,057 | 0,197 | 0,469 | |
| Family | Month | June_vs_September | 0,058 | 0,262 | 0,469 | |
| Family | Month | March_vs_September | 0,03 | 0,501 | 0,501 | |
| **Genus** | **Location** | **SEVo_vs_Ivry** | **0,211** | **0,001** | **0,007** | ** |
| **Genus** | **Location** | **Choisy_vs_SEVi** | **0,379** | **0,001** | **0,007** | ** |

| | | | | | | |
|---|---|---|---|---|---|---|
| **Genus** | **Location** | **Ivry_vs_SEVi** | **0,39** | **0,001** | **0,007** | ** |
| **Genus** | **Location** | **SEVo_vs_Choisy** | **0,243** | **0,002** | **0,008** | ** |
| **Genus** | **Location** | **SEVo_vs_Colombes** | **0,224** | **0,003** | **0,008** | ** |
| **Genus** | **Location** | **Ivry_vs_SECi** | **0,271** | **0,003** | **0,008** | ** |
| **Genus** | **Location** | **Ivry_vs_SECo** | **0,39** | **0,002** | **0,008** | ** |
| **Genus** | **Location** | **Colombes_vs_SEVi** | **0,37** | **0,003** | **0,008** | ** |
| **Genus** | **Location** | **Colombes_vs_SECo** | **0,374** | **0,004** | **0,009** | ** |
| **Genus** | **Location** | **SEVo_vs_SEVi** | **0,238** | **0,005** | **0,01** | * |
| **Genus** | **Location** | **SEVo_vs_SECo** | **0,219** | **0,006** | **0,01** | * |
| **Genus** | **Location** | **Choisy_vs_SECi** | **0,257** | **0,006** | **0,01** | * |
| **Genus** | **Location** | **Choisy_vs_SECo** | **0,393** | **0,005** | **0,01** | * |
| **Genus** | **Location** | **SEVo_vs_SECi** | **0,169** | **0,013** | **0,018** | * |
| **Genus** | **Location** | **Colombes_vs_SECi** | **0,254** | **0,013** | **0,018** | * |
| **Genus** | **Location** | **SEVi_vs_SECo** | **0,147** | **0,014** | **0,018** | * |
| **Genus** | **Location** | **SECi_vs_SECo** | **0,139** | **0,031** | **0,038** | * |
| Genus | Location | SECi_vs_SEVi | 0,077 | 0,662 | 0,772 | |
| Genus | Location | Choisy_vs_Ivry | 0,055 | 0,91 | 0,938 | |
| Genus | Location | Choisy_vs_Colombes | 0,062 | 0,886 | 0,938 | |
| Genus | Location | Ivry_vs_Colombes | 0,052 | 0,938 | 0,938 | |
| **Genus** | **Loc + Month** | **SEVo_vs_Choisy** | **0,58** | **0,001** | **0,002** | ** |
| **Genus** | **Loc + Month** | **SEVo_vs_Colombes** | **0,521** | **0,001** | **0,002** | ** |
| **Genus** | **Loc + Month** | **SEVo_vs_SECo** | **0,561** | **0,001** | **0,002** | ** |
| **Genus** | **Loc + Month** | **Choisy_vs_SECo** | **0,633** | **0,001** | **0,002** | ** |
| **Genus** | **Loc + Month** | **Ivry_vs_SECi** | **0,565** | **0,001** | **0,002** | ** |
| **Genus** | **Loc + Month** | **Ivry_vs_SEVi** | **0,598** | **0,001** | **0,002** | ** |
| **Genus** | **Loc + Month** | **Ivry_vs_SECo** | **0,62** | **0,001** | **0,002** | ** |
| **Genus** | **Loc + Month** | **Colombes_vs_SECi** | **0,544** | **0,001** | **0,002** | ** |
| **Genus** | **Loc + Month** | **SECi_vs_SECo** | **0,491** | **0,001** | **0,002** | ** |
| **Genus** | **Loc + Month** | **SEVi_vs_SECo** | **0,477** | **0,001** | **0,002** | ** |
| **Genus** | **Loc + Month** | **SEVo_vs_Ivry** | **0,546** | **0,002** | **0,004** | ** |
| **Genus** | **Loc + Month** | **Colombes_vs_SEVi** | **0,582** | **0,002** | **0,004** | ** |
| **Genus** | **Loc + Month** | **SEVo_vs_SEVi** | **0,522** | **0,003** | **0,005** | ** |
| **Genus** | **Loc + Month** | **Colombes_vs_SECo** | **0,596** | **0,003** | **0,005** | ** |
| **Genus** | **Loc + Month** | **Choisy_vs_SEVi** | **0,592** | **0,005** | **0,007** | ** |
| **Genus** | **Loc + Month** | **SEVo_vs_SECi** | **0,468** | **0,008** | **0,011** | * |
| **Genus** | **Loc + Month** | **Choisy_vs_SECi** | **0,526** | **0,019** | **0,023** | * |
| **Genus** | **Loc + Month** | **SECi_vs_SEVi** | **0,466** | **0,033** | **0,039** | * |
| Genus | Loc + Month | Choisy_vs_Ivry | 0,461 | 0,055 | 0,061 | . |
| Genus | Loc + Month | Ivry_vs_Colombes | 0,451 | 0,06 | 0,063 | . |
| Genus | Loc + Month | Choisy_vs_Colombes | 0,435 | 0,104 | 0,104 | |
| Genus | Month | January_vs_June | 0,104 | 0,018 | 0,105 | |
| Genus | Month | March_vs_June | 0,082 | 0,035 | 0,105 | |
| Genus | Month | January_vs_March | 0,095 | 0,077 | 0,123 | |
| Genus | Month | January_vs_September | 0,13 | 0,082 | 0,123 | |
| Genus | Month | March_vs_September | 0,084 | 0,125 | 0,145 | |
| Genus | Month | June_vs_September | 0,07 | 0,145 | 0,145 | |
